# Sorption of neuropsychopharmaca in microfluidic materials for in-vitro studies

**DOI:** 10.1101/2021.05.26.445264

**Authors:** Thomas E. Winkler, Anna Herland

## Abstract

Sorption (i.e., ad- & ab-sorption) of small-molecule compounds to polydimethylsiloxane (PDMS) is widely acknowledged. However, studies to date have largely been conducted under atypical conditions for microfluidic applications (lack of perfusion, lack of biological fluids); especially considering the biological studies such as Organs-on-Chips where small-molecule sorption poses the largest concern. Here, we present the first study of small-molecule sorption under relevant conditions for microphysiological systems, focusing on a standard geometry for biological barrier studies that find application in pharmacokinetics. We specifically assess the sorption of a compound panel including 15 neuropsychopharmaca at in-vivo concentration levels. We consider devices constructed from PDMS as well as two material alternatives (off-stoichiometry thiol-ene-epoxy, or tape/polycarbonate laminates). Moreover, we study the much-neglected impact of peristaltic pump tubing, an essential component of the recirculating systems required to achieve in-vivo-like perfusion shear stresses. We find that choice of device material does not significantly impact sorption behavior in our barrier-on-chip-type system. Our PDMS observations in particular suggest that excessive compound sorption observed in prior studies is not sufficiently described by compound hydrophobicity or other suggested predictors. Critically, we show that sorption by peristaltic tubing, including the commonly-utilized PharMed BPT, dominates over device sorption even on an area-normalized basis, let alone at the typically much larger tubing surface areas. Our findings highlight the importance of validating compound dosages in Organ-on-Chip studies, as well as the need for considering tubing materials with equal or higher care than device materials.

## INTRODUCTION

Over the past decades, microfluidic devices and systems have moved from academia into the translational realm.^1^ At the same time, particularly academic applications have introduced more and more biological elements into microfluidic systems, starting with blood for point-of-care testing to complex cell ensembles for recapitulating human organ function in-vitro.^2–4^ Such applications impose ever more rigorous requirements on the materials used in device construction.^5,6^ One major criterium is whether the material alters the behavior or function of the biological element(s) that is to be assessed, i.e. its “biocompatibility”. Thermoplastics (preferred for commercial production) and PDMS (preferred in academic labs) generally perform well on this metric, though a case-by-case assessment is needed. Another important consideration is whether the material interferes with the chemical compounds to be tested, whether talking about a point-of-care biosensor or about drug testing using organs-on-chips.

Studies around the issue of PDMS compound sorption – i.e., non-specific absorption and adsorption, particularly of small molecules – started appearing as early as 2001.^7^ Attention to the phenomenon increased significantly as the microfluidics field continued to expand, especially after a 2006 study considering Nile red and quinine (two small-molecule fluorophores).^8^ It has become accepted that PDMS is capable of absorbing significant quantities of small molecules – especially, though not exclusively, highly hydrophobic ones – into its porous matrix.^9–14^ This distorts not only the absolute compound concentration at steady state (which might be adjusted for, if the uptake percentage is known), but also alters kinetics in time-course experiments, acting as a kind of capacitor – an effect much more difficult (if not impossible) to account for.^15^ The aggregate research on compound sorption has driven studies into limiting such sorption by PDMS surface treatments^7,9,10,16–20^ and into the use of alternate microfluidic materials^12–14,21–23^ – the latter further motivated by PDMS’ other limitations e.g. regarding industrial scale-up.^5,6^

One shortcoming of most, if not all, studies of compound sorption with PDMS and other materials is that they employ simple buffer solutions without proteins/serum. The one exception we have been able to identify found that PDMS sorption was unaffected by the absence/presence of serum, but considered only a single molecule (estradiol).^24^ Protein/serum-free conditions are unlikely to apply to biological sample analysis, and are entirely unsuitable for in-vitro cell culture. Abundant proteins such as serum albumin, however, are likely to adsorb to surfaces and (1) alter their contact angle, e.g. in the case of PDMS rendering it more hydrophilic,^25^ thereby potentially lowering hydrophobic interactions with free hydrophobic compounds; and (2) physically restrict access of small molecules to the surface, potentially decreasing any sorption-related interactions.

Another shortcoming is that microfluidic sorption is considered with a practically exclusive focus on the device itself.^26^ In many scenarios, however, the device represents only a small fraction of the total wetted surface area; liquid reservoirs and especially fluidic tubing (where external pumps are needed) will dominate this by a factor of >10. The assumption of these materials’ irrelevance to study outcomes is perhaps best exemplified by their lack of specification in many Materials & Methods sections. The fact that sorption goes beyond microfluidic materials is implicitly recognized in applied pharmacological studies, where non-specific sorption of the entire system is often measured^27–29^ – but results generally do not translate to other systems.

Herein, we present a study that seeks to assess compound sorption under conditions closely mimicking biological microfluidics, particularly those encountered during in-vitro drug testing in a barrier-on-chip system. We consider 18 small-molecule compounds, mainly neuropsychopharmaca, with a range of physicochemical properties. With these, we characterize material sorption over a typical 24-hour exposure in a complete cell culture media. We prepare microfluidic devices matching a standard design (vertically stacked co-linear channels)^30^ from PDMS as well as two suitable alternatives: (1) off-stoichiometry thiol-ene-epoxy (OSTE+), a polymer system with highly desirable bonding/assembly properties and which features a largely inert and hydrophilic surface after processing;^21^ and (2) thermoplastics in combination with double-coated pressure sensitive adhesive tape (PSA) to facilitate microfluidic device assembly/bonding.^31–33^ Moreover, we compare material sorption of three widely-available peristaltic pump tubings suitable for biomicrofluidic applications:^34^ polypropylene-based (PharMed BPT), silicone-based (Tygon SI), and polyolefin-based (Tygon MHLL).

## DEVICE & STUDY DESIGN

Our microfluidics features dimensions typical for barrier-on-chip research with a cross-section of 1.5×0.2 mm^2^ (*w*×*h*).^35,36^ We follow one of the most established barrier-on-chip designs by placing two such channels on top of each other.^4,15,32,37^ Typically, they would be separated by a porous membrane serving as a cell culture support to facilitate biological barrier formation. To focus on sorption by major structural materials rather than the membrane itself, we instead replace it with a non-porous polycarbonate (PC) film. The resulting devices and flow paths, which enable us to utilize just one pump channel per device, are illustrated in Figure 1.

**Figure 1.**
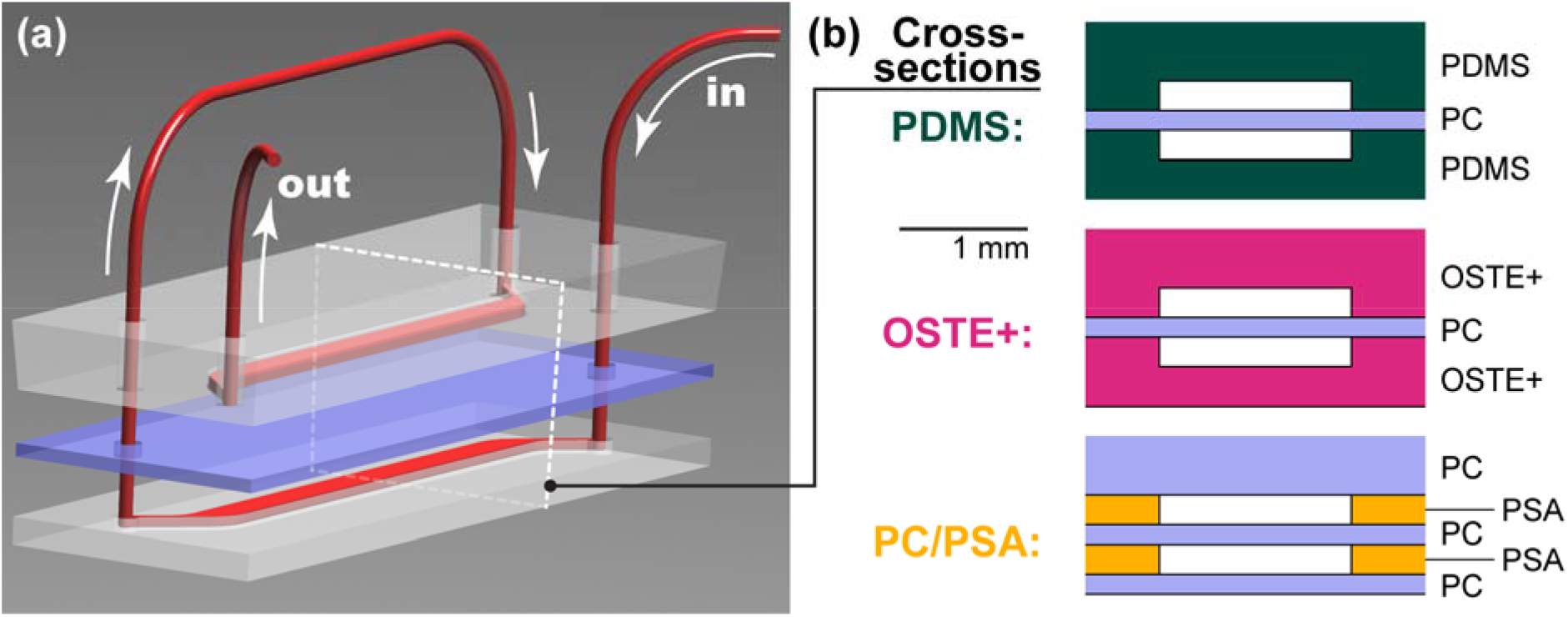
Barrier-on-chip-type microfluidic devices. **(a)** Schematic 3D rendering, showing the media perfusion (red) through the microfluidics from the **(in)**let to the **(out)**let. The internal wetted microfluidic area is 1 cm^2^. White dotted line indicates cross-section plane. **(b)** Cross-sections for the three material options we consider in our study. Abbreviations: PDMS – poly(dimethylsiloxane); OSTE+ – off-stoichiometry thiol-ene-epoxy; PC – polycarbonate; PSA – pressure-sensitive adhesive.

The epithelial or endothelial cells typically employed in barrier-on-chip devices can greatly benefit from physiologically relevant shear (τ = 1∼10 dyne cm^−2^).^38–40^ To achieve similar forces in microfluidic geometries, we consider the equation (suitable for *w*>*h*):

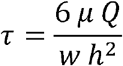

With geometries in the hundreds of microns, this means that flow rates *Q* in the range of milliliters per hour are required, even when adjusting viscosity μ to in-vivo-like values.^41^ With a multiplexed, multi-day experiment, this adds up to an unsustainable volume of media (and associated amounts of drugs or similar test compounds), thus necessitating media recirculation. These requirements (recirculation & multiplexing) make peristaltic pumps the obvious (and in most cases, only) answer.^42^ Only a limited variety of tubing materials is available, however;^34^ the selection is even more limited when requiring biocompatibility as per USP Class VI or ISO 10993 (reasonable minimum standards for in-vitro applications). This leaves us with three classes of materials (we use market-dominant Tygon plastics; alternative manufacturers’ products will still fall into the same categories): polypropylene-based (PharMed BPT), silicone-based (Tygon SI), and polyolefin-based (Tygon MHLL). PVC-based tubing (e.g. Tygon LFL) may be USP-certified, but is not recommended for usage with blood/tissue due to plasticizer leaching concerns.

As sorption test compounds, we select 3 fluorophores and 15 neuropsychopharmaca (Table 1). Brain-targeting drugs with their generally high hydrophobicity (to ease passage across the blood-brain-barrier)^43^ present a particularly difficult scenario. The specific group of compounds was selected based on (i) their known clinical need for therapeutic drug monitoring (TDM),^44^ as this can indicate the relevance of in-vitro time-course studies as well as the desirability of labs-on-chips for point-of-care analysis; (ii) their spread in hydrophobicity to encompass a range of properties;^45^ and (iii) the non-restricted availability of the compounds themselves, as well as of standardized clinical LC/MS analysis. Overall, their properties are representative of brain-targeting drugs in general (mean *M* ∼ 310 Da, H-Bd < 3), with a bias towards higher hydrophobicity (literature median log*P*: 2.5; ours: 3.7) to increase the likelihood of observing material sorption.^46^ Combined with concentrations *C* at 5/6^th^ of the upper therapeutic range limit in plasma,^44^ our compound panel represents realistic conditions for pharmacokinetic studies in a barrier-on-chip system.

**Table 1.**
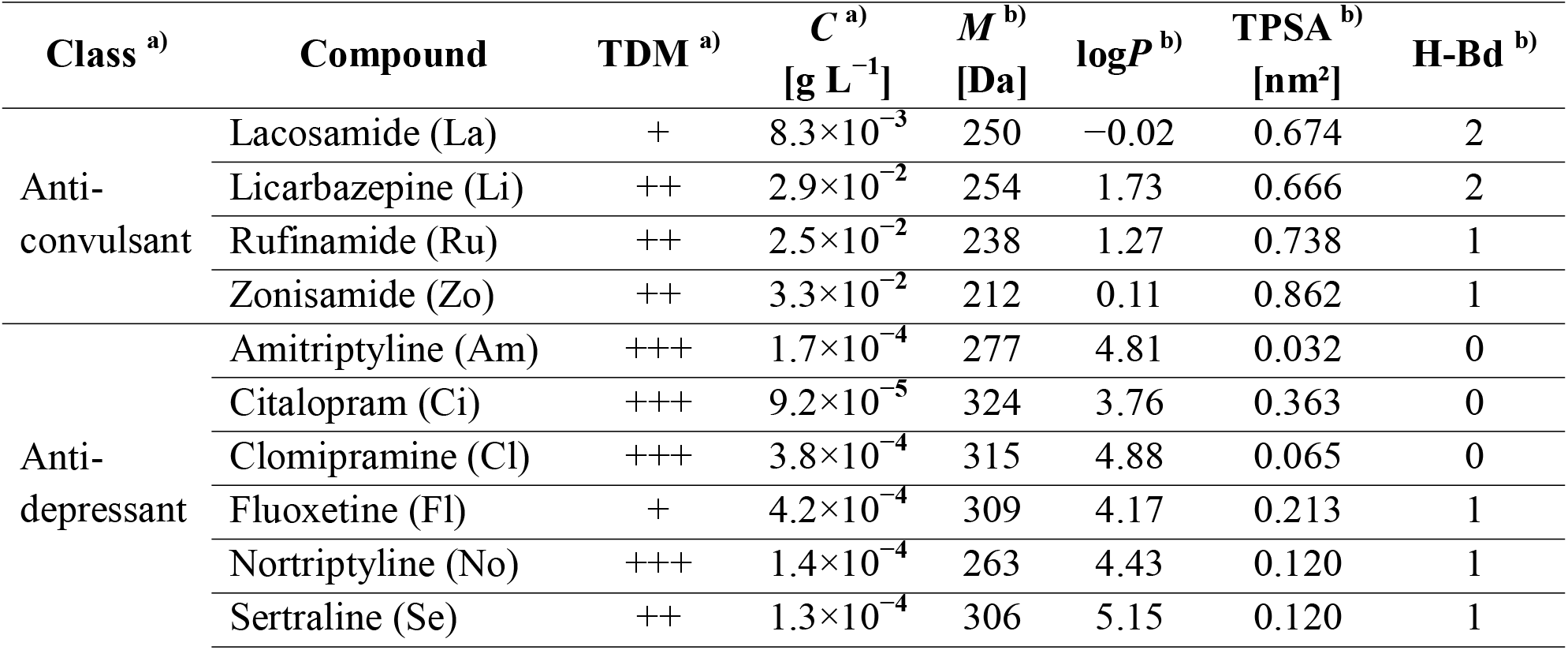
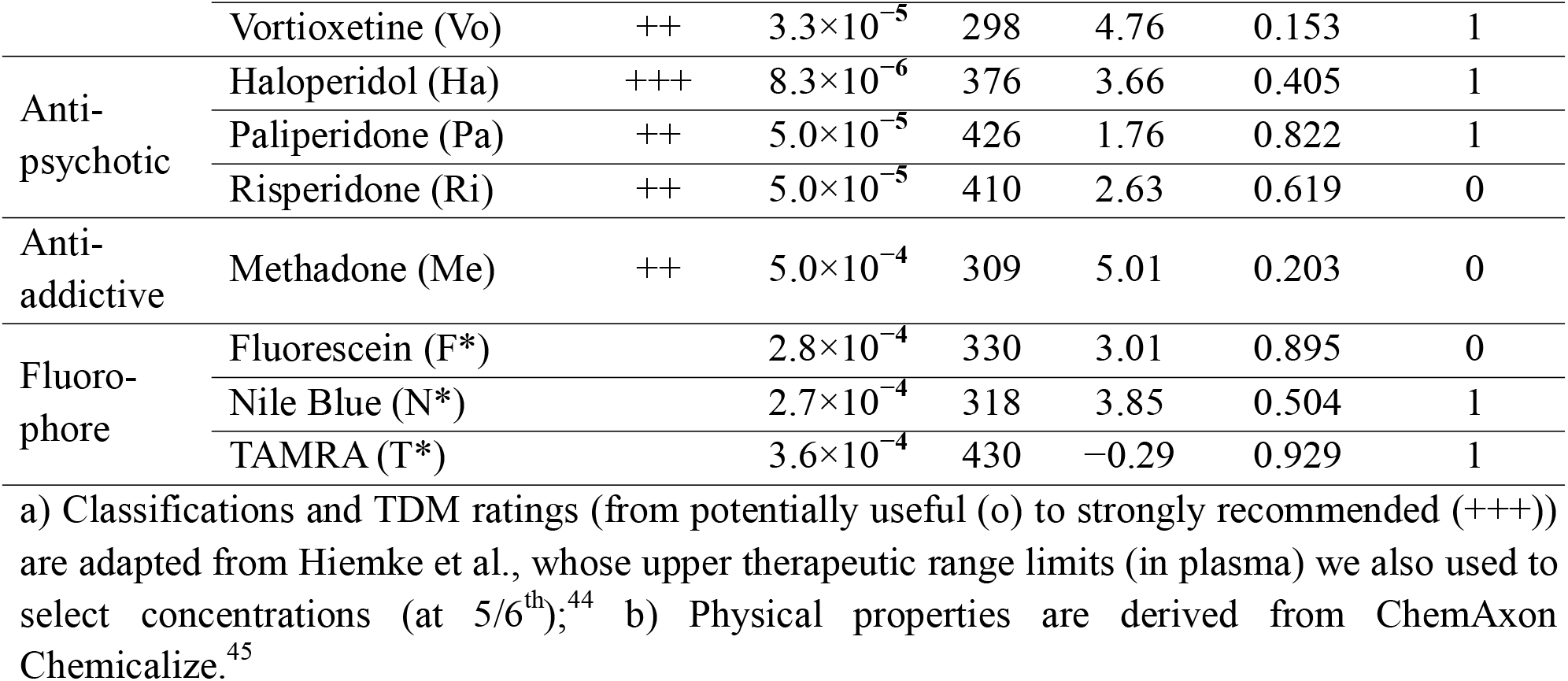
Compounds selected for study, as well as relevant physical and chemical properties. TDM, therapeutic drug monitoring; *C*, concentration in our study; *M*, molar mass; *P*, octanol-water partition coefficient; TPSA, topological polar surface area; H-Bd, H-bond donor count; TAMRA, tetramethylrhodamine

## MATERIALS & METHODS

### Exhaustive descriptions are provided as Supplementary Information

As mentioned in the previous Section, all our devices featured identical channel geometries with an internal wetted surface area of 1 cm^2^. Overall (external) chip design and dimensions were adjusted to accommodate the various fabrication processes. PDMS (∼2.6 ml/device) was cast on photolithographically structured molds,^47^ and devices sealed by external clamping. We fabricated OSTE+ devices using UV-reaction-injection molding followed by thermal bonding.^48–50^

For PC/tape structures, we relied on knife-plotter cutting and pressure-assisted lamination.^32^ We substituted solid PC films for the typical porous membranes (often PC, PET, or PDMS) in all process flows. PharMed BPT tubing was used for all connections, including to guide the fluid as shown in Figure 1. Besides devices, the total wetted surface area overall included: 7.1 cm^2^ tubing (including peristaltic pump tubing); 0.8 cm^2^ stainless steel interconnects; ∼7 cm^2^ fluid reservoirs (at typical fill volume). For peristaltic pump tubing evaluation (PharMed BPT, Tygon SI, Tygon MHLL), we constructed flow circuits of equal length, omitting only the microfluidic devices.

On day −1, we plasma-treated all devices and tubing to facilitate wetting. After connection to the peristaltic pump, everything was flushed (i.e., liquid displacement >5× total internal volume) with 70% ethanol over 10 minutes for disinfection and moved into a standard cell culture incubator. We continually perfused devices with PBS overnight to calibrate flow speeds and check for leakage. On day 0, we flushed the system with cell culture media (with 10% serum replacement, except where we note supplement-free media (SFM)) and set up recirculating flow by looping pump outlets back into the liquid reservoirs. For recirculating flow, we utilized a total media volume of 2.5 ml and a linear pump speed of 17.5 mm s^−1^, corresponding to *Q* = 4 ml h^−1^ and τ = 1.1 dyne cm^−2^ (cf. Equation 1) with devices. On day 1, we flushed everything with fresh media spiked with pharmaceuticals and dyes according to Table 1 before continuing recirculating flow. After another 24 h (day 2), we collected 1 ml samples for analysis from each of the reservoirs.

Fluorophore analysis was performed with a plate reader. We sent samples for standard TDM analysis at the local hospital’s clinical laboratory facilities. We imaged tubing cross-sections using a tabletop SEM. For data analysis, we relied on Origin Pro (statics, plots), MOVER-R^51^ (confidence interval ratios), and Fiji^52^ (micrographs).

## RESULTS & DISCUSSION

### Microfluidic Devices

First, we consider compound sorption for our microfluidic devices. To extract only the device contribution, we compare all results to corresponding tubing-only experiments (i.e., identical fluidic circuits omitting the devices). This corrects for device material-independent losses such thermal degradation (Figure S1) or sorption by other parts of the setup (discussed in a later Section; Figure 3). The resulting fractional compound recovery for microfluidics from various materials is shown in Figure 2a. Along the *x*-axis, we sort compounds by their topological polar surface area (TPSA), a measure of hydrophilicity (lower TPSA = more hydrophobic). The choice is based on van Meer et al.’s finding wherein PDMS sorption correlated better with TPSA than with the octanol-water partition coefficient log*P*, the more commonly-used hydrophobicity metric.^10^ Another advantage of TPSA is that calculations are less divergent between prediction algorithms (compared to log*P*). Of the 13 compounds detected (cf. next Section), we find that – independent of device material – all of their 95% confidence intervals envelop a recovery efficiency of 1.0 (i.e., zero compound loss/sorption). The per-compound confidence intervals also strongly overlap between material conditions. We note however a decrease in means toward the left side of the plot, suggesting potential trends when considering all compounds. A linear regression analysis confirms that sorption noticeably increases with increasing hydrophobicity for PDMS as well as OSTE+ (but not PC/PSA; the relevant data are included as part of Table 3 in the next Section).

**Figure 2.**
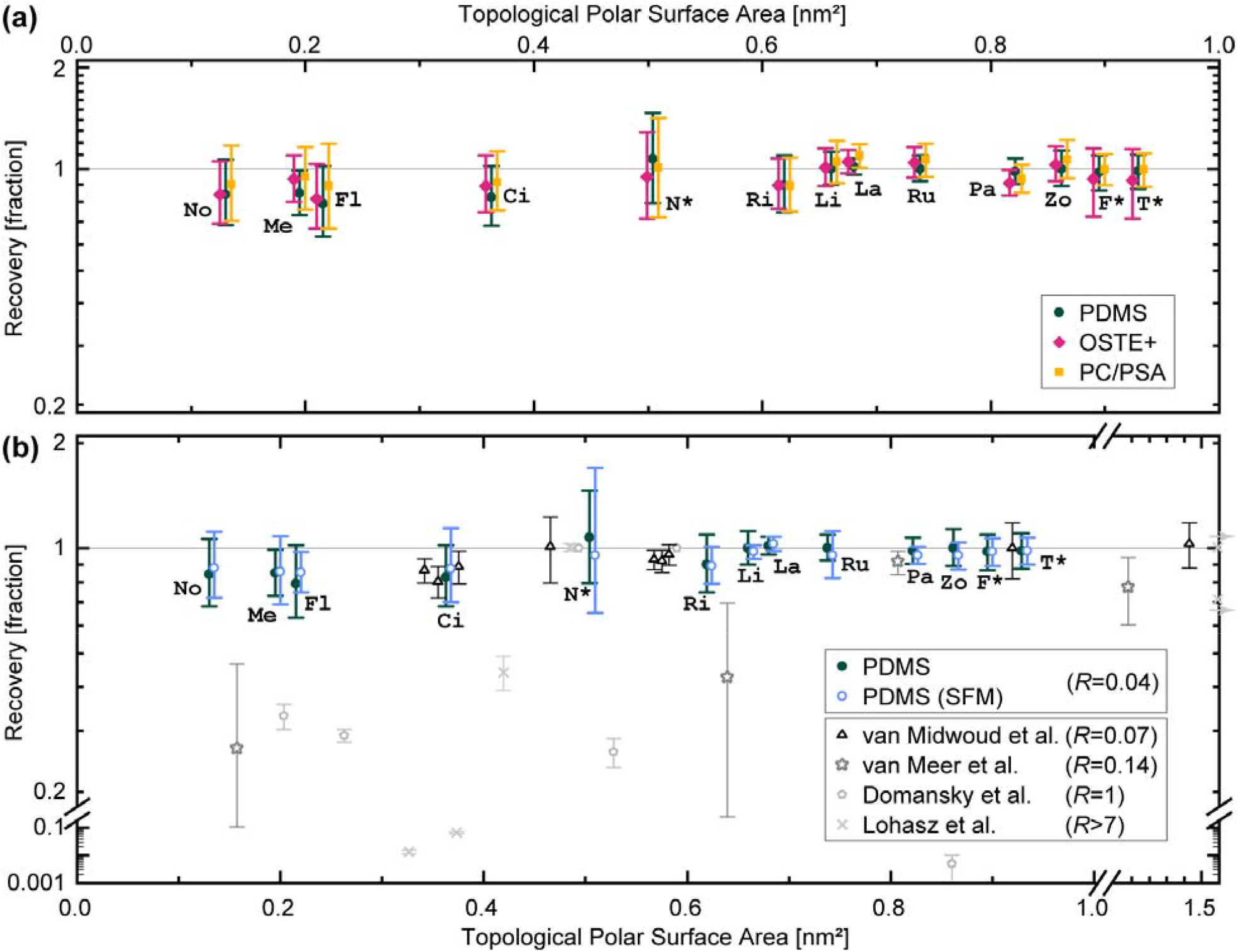
Sorption in microfluidic devices. Compound recovery from microfluidic devices after 24 h of continuous recirculating media perfusion, normalized to tubing-only controls. **(a)** We compare device materials as listed in the legend and illustrated in Figure 1. **(b)** Comparison of PDMS results for media with and without proteins/serum (SFM: supplement-free; otherwise 10% serum replacement). Data from four other studies (featuring suitable control data for necessary corrections; see text) are included for reference.^10,12–14^ All compounds are sorted by their hydrophobicity in terms of topological polar surface area (offset for better visualization where needed); for abbreviations see Table 1. Data are plotted as means (*n*=4 per condition), with error bars representing the 95% confidence interval. For literature values, error bars represent SD (*n*=3)^12,14^ or range (*n*=2),^10,13^ and arrows on right border indicate protein-size compounds.

**Figure 3.**
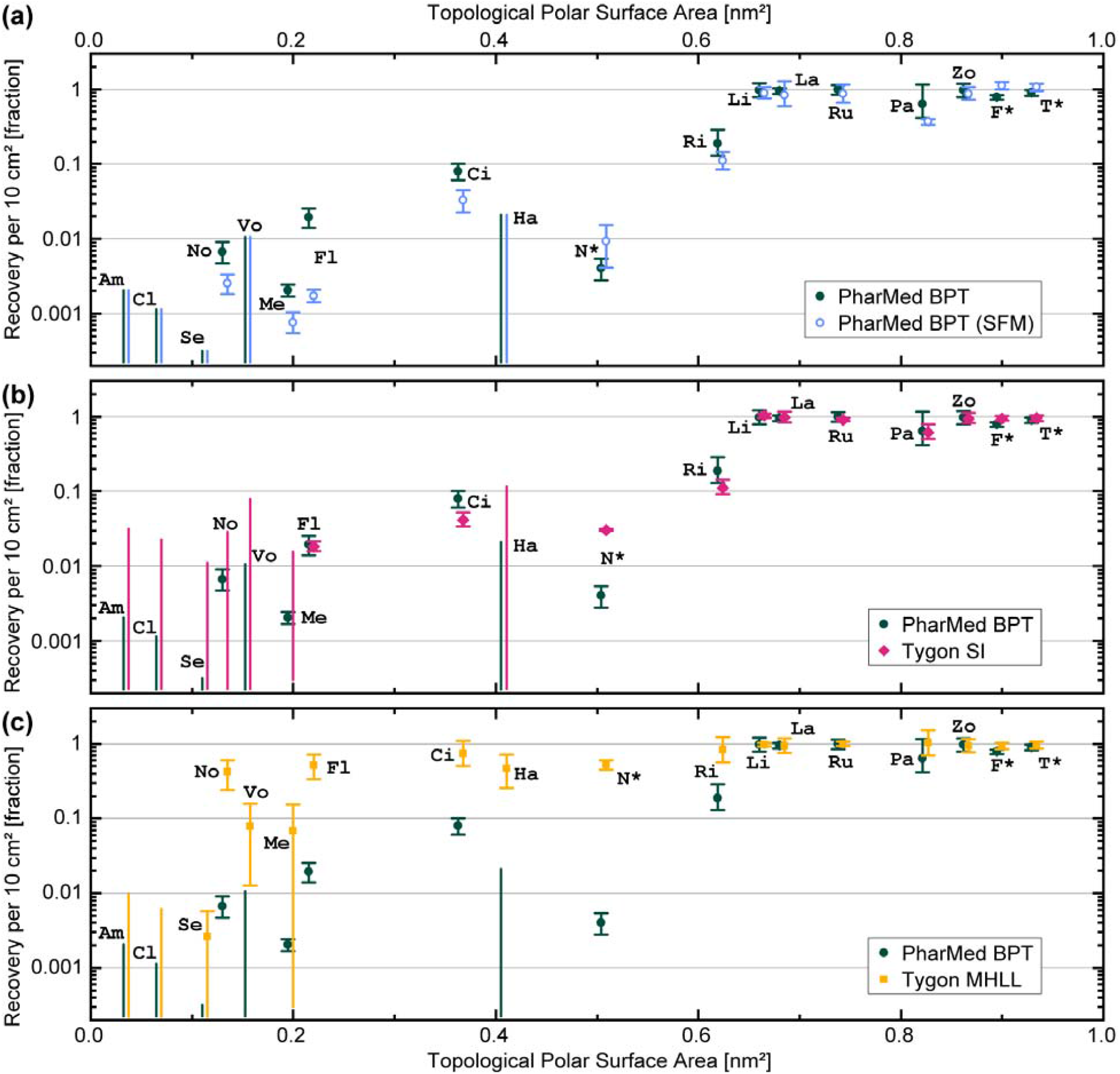
Sorption in microfluidic tubing. Compound recovery (as a fraction) from tubing-only flow circuits after 24 h of continuous recirculating media perfusion (10% serum). The graphs compare PharMed tubing with **(a)** same tubing, but serum-free media; **(b)** silicone-based tubing; and **(c)** MHLL-type tubing. All values are normalized to 10 cm^2^ tubing area to highlight the typical surface area mismatch with devices. Neuropsychopharmaca are sorted by their hydrophobicity in terms of topological polar surface area (offset for better visualization where needed); for abbreviations see Table 1. Data are plotted as means (*n*=3 per condition), with error bars representing the 95% confidence interval. Uncapped error bars correspond to the below-LOQ range for conditions where data < LOQ.

**Table 2.**
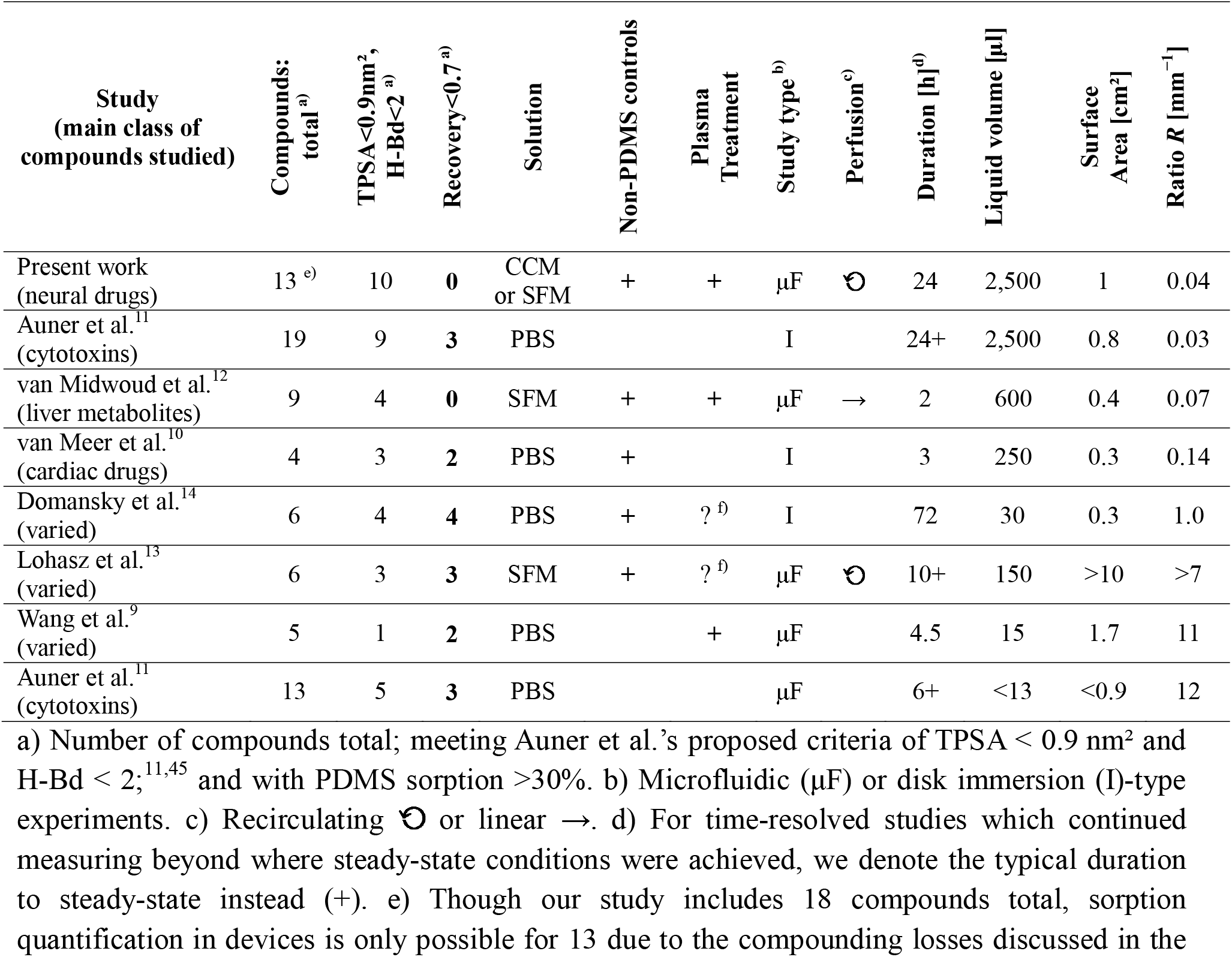
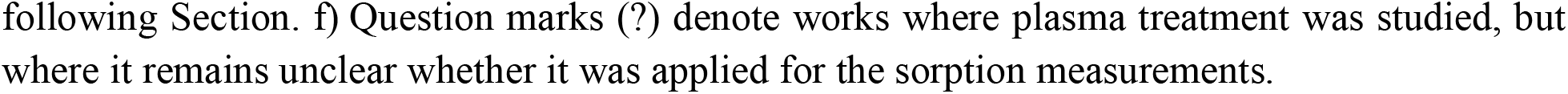
Overview of studies on PDMS compound sorption. The table summarizes some of the relevant parameters of the studies. We exclude studies with <3 compounds. Values are estimated from available data where not explicitly stated in the relevant papers. *R*, surface area-to-liquid volume ratio.

**Table 3.** Dependence of compound sorption on TPSA for the various device and tubing materials considered in our study. For comparison purposes, all sorption values are normalized to 1 cm^2^ material surface area (i.e., the device equivalent)

| **Material** | **Slope <sup>a)</sup>** | **(95% CI) <sup>a)</sup>** | **Pearson's *r*** |
| --- | --- | --- | --- |
| **Devices** |  |  |  |
| PDMS | **0.08** | (0.04; 0.13) | 0.76 |
| PDMS (SFM) | **0.07** | (0.03; 0.12) | 0.71 |
| OSTE+ | **0.06** | (0.00; 0.13) | 0.56 |
| PC/PSA | **0.02** | (−0.06; 0.09) | 0.14 |
| **Tubings** |  |  |  |
| PharMed BPT | **0.28** | (0.20; 0.36) | 0.88 |
| PharMed BPT (SFM) | **0.36** | (0.30; 0.42) | 0.95 |
| Tygon SI | 0.42 <sup>b)</sup> | (0.34; 0.50) <sup>b)</sup> | 0.94 |
| Tygon MHLL | **0.05** | (0.01; 0.08) | 0.58 |

a) Slope, and corresponding confidence interval CI, in terms of log(recovery) / log(TPSA). b) Tygon SI data fitting is unreliable due to the unusually narrow confidence interval in its N\* measurement ( $\pm 0.002$ , vs. around $\pm 0.02$ for other materials), and a strong dependence of the slope on this particular data point.

Putting the PC/PSA result in context is difficult, as ours appears to be the first quantitative PSA sorption study. The one extant study we were able to locate on PSA sorption only provides qualitative analysis for a single fluorescent dye.^53^ They observe limited sorption of dye, especially in the presence of proteins, under ambient conditions. This is very broadly in agreement with our own findings. In our study, sorption of PSA is at the very least not high enough to outweigh the benefits of a thermoplastic (PC) constituting the majority of the microfluidic surface area. In one comparative study, PC sorption was generally on par with polystyrene (the most common cell culture plastic), and behind only cyclic olefin copolymer (COC); all three significantly better than PDMS.^12^ Based on our trend analysis, PC/PSA may be considered a potentially optimal choice, even if the confidence intervals largely overlap the other conditions.

OSTE+ sorption has previously been studied qualitatively and quantitatively, but only with rhodamine B in water.^54–56^ These findings generally showed that OSTE+, unlike PDMS, does not exhibit absorption (as expected for a tightly crosslinked matrix), but that it does feature noticeable surface adsorption. This still presents a significant potential advantage, since surface sites can saturate and also be modified with appropriate surface chemistry. PDMS devices, conversely, can continue to absorb compounds into loosely-crosslinked internal volumes many times that of the microfluidic channels they contain. Our own results for OSTE+ cannot establish whether it has any absolute advantages over PDMS regarding compound sorption. However, based on the hydrophobicity trend analysis, it performs at least similarly to PDMS (which itself performs well), or better.

PDMS allows for the most expansive discussion, due to a larger existing body of research. In our study, the lower ends of the confidence intervals all remain above 0.6. Our findings do indicate that hydrophobicity impacts the sorption behavior of PDMS, but only with a small effect size, and without any outliers. This is a significant contrast to prior studies (cf. Table 2), which generally report sorption in excess of 0.5 log_10_ (∼30%) for at least half of their tested compounds below a proposed TPSA threshold for sorption of 0.9 nm^2^.^9–11,13^ To explain why some (but not all) compounds below the cutoff exhibited high sorption, Auner et al. consider 15 chemical and physical descriptors.^11^ They suggest that, below the TPSA cutoff, the number of H-bond donors H-Bd – potentially related to molecular diffusivity within PDMS – becomes the determining factor (high sorption if 0; also more likely to exhibit high sorption at 1, and potentially at 2). However, as per Table 1, ten of our measured compounds here fall below the TPSA cutoff and satisfy H-Bd < 2 (Me, Ci, and Ri at 0). This suggests that the H-bd criterion may have arisen from compound selection bias. It is worth noting that Auner et al.’s study (the only one to exceed ours in compound number) focused on toxins rather than (as in our study) pharmaceuticals, covering a range also of more hydrophilic molecules, as well as compounds with H-Bd > 2.^11^ following Section. f) Question marks (?) denote works where plasma treatment was studied, but where it remains unclear whether it was applied for the sorption measurements.

For additional context, we first consider the various time/volume/surface parameters from prior research. We would expect shorter exposure duration, and lower PDMS surface-to-liquid volume ratios *R*, to correlate with lower sorption. *R* clearly divides the studies in Table 2 into two groups (Domansky et al.’s work notwithstanding). Those with *R*∼10 mm^−1^ would all be expected to show higher sorption based on this fact alone, and comparison to those with 100-fold lower *R*∼0.1 mm^−1^ should almost certainly be avoided. Indeed, the fraction of high-sorption compounds is higher in the latter group (Table 3). Even within the low-*R* group, to which our study belongs, however, reported differences in sorption behavior are sizable. We will attempt to address some of the potential confounding factors in the following paragraphs: sample solution type, plasma treatment, inclusion of non-PDMS controls, and finally a closer look at time and *R*.

One difference of our study compared to prior research is the, for in-vitro modeling realistic, use of complete cell culture media (CCM; rather than protein/serum-free media (SFM), or buffer solutions (PBS)). Since we include both CCM and SFM conditions (Figure 2b), with SFM exhibiting practically identical trends, we can however rule out protein content as a dominant factor in PDMS compound sorption. This refutes one of our initial hypotheses, and provides supporting evidence for this previously estradiol-only finding.^24^ In terms of sample solutions, Table 2 also shows that studies finding high sorption in the low-*R* group rely on PBS rather than even on SFM. However, we conclude that this is not a significant factor due to the lack of a similar correlation in the high-*R* group (combined with the generally hydrophilic nature of SFM constituents, which should not interfere with hydrophobic sorption).^57^

Auner et al. speculate that plasma treatment could play a role in PDMS compound sorption.^11^ Indeed many extant studies (also beyond those in Table 2) consider untreated PDMS even in microfluidic test formats,^8,11,13,24,58^ whereas we include this as a step typically performed prior to realistic use. Wang et al.’s observation of up to 80–95% losses in their plasma-treated channels – with otherwise nearly identical study design to Auner et al.’s microfluidic experiments – is a strong argument against plasma treatment as a dominant factor.^9^

We find a more important caveat in that prior studies (excepting Domansky et al.)^14^ report compound loss relative to starting concentrations.^9–13^ Potential losses from inherent compound degradation (due to time, temperature, *p*O_2_, …) are thus conflated with PDMS sorption. In two studies (Lohasz et al., van Midwoud et al.), control data with thermoplastic materials can establish that such losses were negligible;^12,13^ no such data are provided by Auner et al. (closest to our study in *R*), and in van Meer et al.’s experiments, comparison to those control values (rather than initial) would decrease the reported PDMS sorption figures up to half.^10^ The data displayed in Figure 2b include the appropriate corrections.

Considering all relevant parameters now, our study is most similar to the one by van Midwoud et al., and indeed theirs is the only other multi-compound study finding PDMS sorption in a range (0–20%; Figure 2b) similar to our own data.^12^ Although our own study lags behind van Midwoud et al.’s in terms of *R* by a factor of 2, based on our longer liquid residence times (duration and recirculation) we may still have expected a somewhat larger effect size than their study. Van Meer et al.’s study also points to a larger expected effect size (even accounting for inherent losses mentioned above), since their larger *R* should be compensated for by our longer duration.^10^ The nature of the experiment (disk immersion) is naturally different, but diffusion (∼1 cm/24 h) limits the PDMS-exposed molecules per unit time similarly as perfusion does in microchannel experiments like ours.

Our PDMS data, and the above discussion, point to PDMS sorption being strongly dependent on *R* as well as on the specific compounds chosen. PDMS clearly exhibits compound sorption behavior based on hydrophobicity in terms of TPSA. However, the effect size is – for *R* in line with a typical in-vitro study – smaller than generally assumed. Exceedingly high-sorption compounds (>30%) appear to be outliers, rather than the norm, even when only considering hydrophobic ones (as evidenced also by the lack of clear TPSA-dependence in the Domansky et al. and Lohasz et al. data in Figure 3b). None of the criteria suggested to date seems sufficient to predict these with certainty. Overall, our data suggest that – for a barrier-on-chip-type experiment – compound sorption is limited, and not significantly impacted by the choice of device material.

### Microfluidic Tubing

As we mentioned in the Introduction and Study Design Sections, the microfluidic device is often only a small part of the entire set-up. For the barrier-on-chip scenario we are modeling here, the tubing itself makes up most of the exposed fluidic area (∼7 cm^2^; the reservoirs contribute a similarly-sized area, but allow for use of low-sorbing plastics like polypropylene in our case).^59^ To analyze the tubing contribution, we now consider the compound recovery from tubing-only fluidic circuits (which served as controls in the prior Section) relative to samples exposed to the same environment in a low-sorption plastic vessel. The latter (when compared to initial concentrations) yield information on inherent degradation from environmental factors, which is appreciably non-zero for three compounds (between 25 and 50% for Am, Cl, Se; Figure S1).

In Figure 3, we plot the compound sorption from three tubing materials: polypropylene-based (PharMed BPT; 3a), silicone-based (Tygon SI; 3b), and polyolefin-based (Tygon MHLL; 3c). We find that all three tubings show significant sorption of at least half our study compounds, well in excess of our microfluidic devices. Some compound concentrations decrease enough to fall below our assay limit of quantitation, which is also indicated in the graphs (the affected compounds therefore not possible to include in the earlier device analysis). We observe that sorption under all conditions depends strongly on compound hydrophobicity in terms of TPSA. Indeed, multivariate partial least squares models confirm that log(TPSA) is the most important molecular parameter across all materials (score 1.9±0.1). Out of 21 chemical properties, plus the experimental concentration parameter log(*C/M*), only the other hydrophobicity measures log*P* (1.7) and log*D*_7.4_ (1.5) score significantly higher than 1 on variable importance, as does the H-bond <u>acceptor</u> count (1.4). As examples, we plot the correlations with log*P*, log(*C/M*), and molar mass *M* in Figure S2 (see Supplement for full list of model parameters).

We can further quantify the sorption trend regarding TPSA using a linear regression model (in log-log space; showing highest linearity). For the resulting data in Table 3, we normalized to a device-equivalent area of 1 cm^2^ for direct comparison with the prior Section. These values show that not only is compound sorption higher than device sorption due to larger tubing surface areas (as represented in Figure 2 versus Figure 3); tubing material sorption can be much higher even on an equivalent-area basis. Thus tubing materials, and choice of tubing, are much more critical than device material choices for in-vitro microphysiological studies of the type we mimic here. We will discuss the various materials in further detail in the following paragraphs.

PharMed BPT (polypropylene-based) is a widely used option for in-vitro models both by us and other groups,^4,15,32,60–64^ and thus serves as our default. In Figure 3a, we first consider sorption of this material for both complete cell culture media as well as protein/serum-free media (SFM). With SFM, five individual compounds show significantly reduced recovery (in terms of non-overlapping confidence intervals), and the aggregate slope parameter for PharMed BPT increases (albeit retaining overlapping confidence intervals; Table 3). Thus, unlike with our earlier observations on PDMS, the presence of proteins does appear to passivate the surface regarding sorption for this material. In either case, PharMed BPT sorption is clearly non-negligible. This is in spite of its parent plastic (polypropylene) generally being considered low-sorption;^59^ the thermoplastic extrusion process to create a flexible material, and the plasticizers used, must thus play a role in the observed behavior. SEM on tubing cross-sections (Figure 4a) provides supporting evidence for the plastic processing hypothesis. Compared to the other materials, PharMed BPT features a sponge-like structure, also exhibiting a highly textured liquid-facing surface. At the very least, this increases the effective surface area of the material significantly beyond what the tubing dimensions alone imply. We note that one prior study by Chao et al. did characterize sorption in their entire in-vitro model system with PharMed BPT (using SFM) compared to a highly specialized alternative (no longer on the market).^29^ Considering 4 compounds, they reported high PharMed BPT sorption (>90%) for 2 of them; the lack of geometric parameters (diameter, length) unfortunately does not allow for extrapolation or comparison beyond their system.

**Figure 4.**
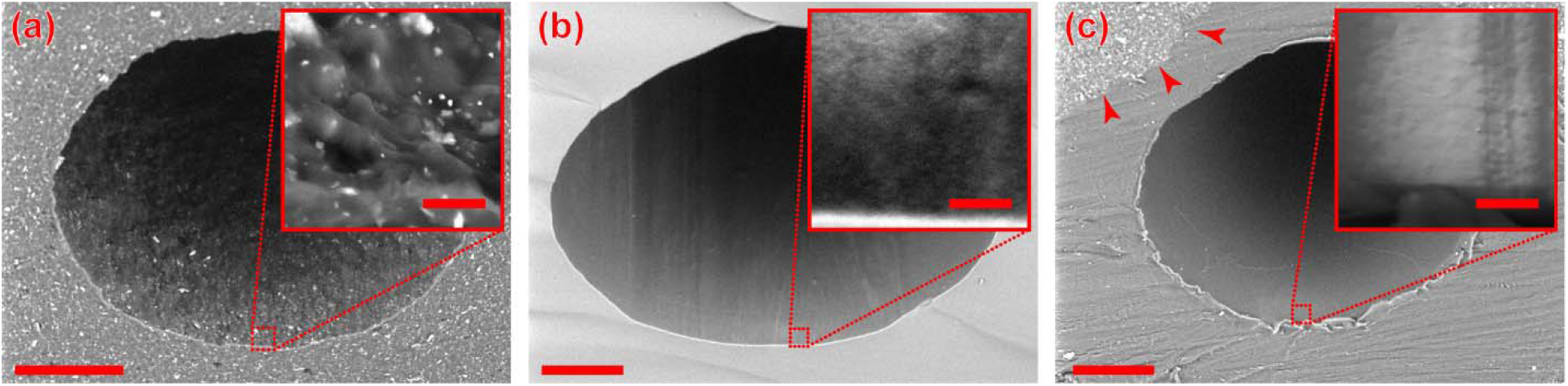
SEM images of tubing materials. Cross-section micrographs – with insets showing high-magnification views of the inner tubing surface – for (a) PharMed BPT; (b) Tygon SI; (c) Tygon MHLL. With Tygon MHLL, the arrows indicate where the inner polyolefin (Tygon MH) core transitions into the outer PharMed BPT sheath. Scale bars are 100 μm (insets: 5 μm).

Tygon SI is also utilized in the context of microphysiological models.^65–67^ Intuitively, based on the underlying plastics involved (silicone versus polypropylene), this should be a higher-sorbing tubing, a reasoning specifically mentioned e.g. by Bovard et al. in their choice of PharMed BPT.^64^ In our study, as per Figure 3b, interestingly only one compound (Ci) exhibits significantly lower recovery with Tygon SI, whereas another (N*) actually shows the reverse behavior. We hypothesize that the difference in surface roughness plays a role here, with Tygon SI featuring a very smooth interior surface (Figure 4b). Compared to the highly textured surface of PharMed BPT, this can compensate for a presumed higher inherent silicone sorption. Based on our results, other practical differences should therefore be used as the main decision factors between PharMed BPT, such as pump life (PharMed BPT), transparency (Tygon SI), or oxygen permeability (Tygon SI).^34^

Tygon MHLL consists of an inner layer of high-purity polyolefin-based plastic (Tygon MH), coated with an outer shell of PharMed BPT to increase flexibility for peristaltic pump use (Figure 4c). We were unable to identify any in-vitro model studies employing this tubing, potentially due to its high stiffness necessitating manual peristaltic pump adjustments when accurate flow control is desired.^34^ When comparing against PharMed BPT tubing in Figure 3c, we find that Tygon MHLL performs markedly better. Recovery is significantly improved for 9 compounds, and the aggregate analysis reveals a much-flattened slope (Table 3). The result is broadly in agreement with Chao et al., whose aforementioned study also compared Tygon MHLL to PharMed BPT for a single compound, finding 20-fold improved recovery.^29^ Even this material, however, still suffers from compound sorption of greater than 99% for our 3 most hydrophobic compounds (Am, Cl, Se; at 10 cm^2^ surface area). Lacking a better commercially-available alternative, our data suggest Tygon MHLL is nonetheless the optimal peristaltic pump tubing for in-vitro modeling with highly hydrophobic pharmaceutica.

## CONCLUSION

We sought to characterize sorption in microfluidic systems – uniquely focusing on *both* devices and requisite tubing – for in-vitro studies under realistic conditions. To that end, we employed one of the largest and most hydrophobic compound panels to date, mainly consisting of neuropsychopharmaca. In contrast to many prior reports on PDMS, we did not find any compounds exhibiting 2+ fold-changes in concentration from device materials alone, attributing this in large part to discrepancies in surface-to-volume ratios. Comparing PDMS to two alternative device construction methods – OSTE+ or PC/PSA – showed that these perform at least on par if not better than PDMS. We further found that sorption of tubing materials like the widely-used PharMed BPT has a strong dependence on compound hydrophobicity, and critically dominates over device material sorption even on an area-normalized basis (let alone with typically ∼10-fold larger tubing areas). Tygon MHLL performed best among the readily-available pump tubing options we evaluated.

We acknowledge that, in our combined study design, the high level of tubing sorption likely lowered our assay sensitivity for device materials alone. Thus, we hope to verify potential advantages of PC/PSA in future experiments with more optimized tubing circuits (see below). Regardless, our results suggest that orders-of-magnitude sorption reported with PDMS represents compound outliers more than the rule, and that prior suggested predictors are at most necessary but clearly not sufficient. We believe that computational approaches, including more advanced machine learning to identify relevant molecular properties (as applied to prediction of e.g. blood-brain barrier permeability),^68^ as well as molecular dynamics modeling,^69^ present logical next steps towards understanding this behavior.

Overall, we see the implications of our research for biomicrofluidics – especially toward in-vitro pharmacodynamic modeling – as three-fold: First, confirmation of compound concentrations, as well as inclusion of proper controls, is necessary. Second, all fluidic components – beyond device materials alone – need to be specified in Methods sections. Third, “soft” peristaltic pump-type tubing should be minimized as much as possible, replaced with glass-lined or fluoropolymer-based tubings. Although this is widely acknowledged in the microfluidics community, the downsides of flexible tubing had not been highlighted this clearly to date. The advantages of peristaltic pumps are too great to fully avoid them, but the relevant tubing can be optimized (Tygon MHLL or similar) and shortened; it also provides an incentive toward micropump integration.^70^ Ultimately, tubing and associated materials clearly deserve at least the same amount of attention as device materials, if not more.

## AUTHOR INFORMATION

## Author Contributions

**T.E.W**.: Conceptualization, Methodology, Investigation, Analysis, Writing, Administration;

**A.H**.: Conceptualization, Review & Editing, Administration, Supervision

## ACKNOWLEDGMENT

We appreciate the assistance of the Karolinska University Hospital Centre for Clinical Laboratory Studies. We further appreciate the support of Mikael Bergqvist (KTH Micro- and Nanosystems) as well as Laurent Barbe (SciLifeLab Customized Microfluidics) in device fabrication.

## ABBREVIATIONS

PDMS: poly(dimethylsiloxane)
DMSO: dimethylsulfoxide
PBS: phosphate buffered saline
LC/MS: liquid chromatography/mass spectrometry
LOQ: limit of quantitation
PC: polycarbonate
MEM: minimum essential medium
CCM: complete cell culture media
SFM: supplement-free media
TDM: therapeutic drug monitoring
TPSA: topological polar surface area
TAMRA: carboxytetramethylrhodamine
*R*: surface area-to-liquid volume ratio
SEM: scanning electron microscope
CI: confidence interval
SD: standard deviation.

