## Supporting Information for "Sorption of neuropsychopharmaca in microfluidic materials for in-vitro studies"

### Supporting Information:
### Sorption of neuropsychopharmaca in microfluidic materials for in-vitro studies

*Thomas E. Winkler[1]* and Anna Herland[1,2]**

[1] Division of Micro- and Nanosystems, KTH Royal Institute of Technology, 10044 Stockholm, Sweden.

[2] AIMES, Department of Neuroscience, Karolinska Institute, Solna, Sweden.

*

NB: The supplemental Detailed Experimental Section is located after the supplemental Figures & Tables referred to in the main manuscript.

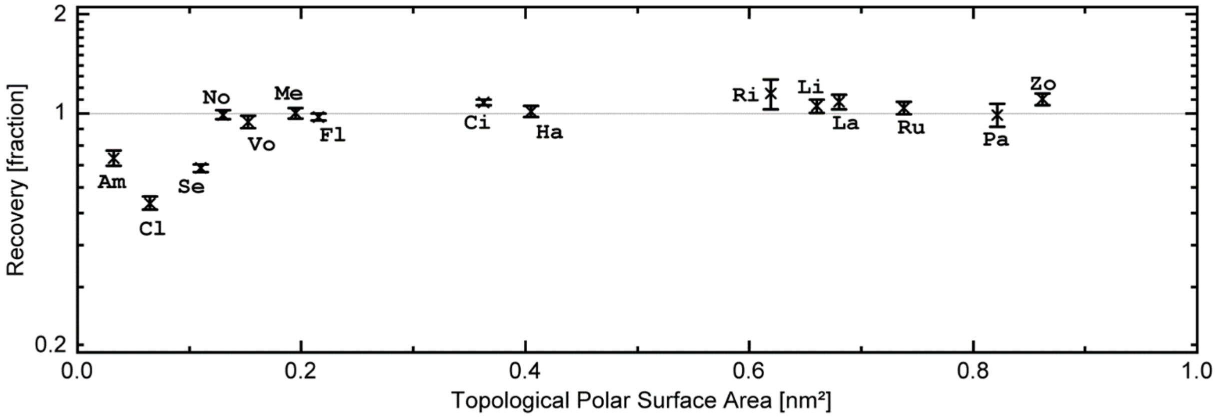


**Figure S1.** Thermal/environmental degradation and loss of compounds. Drug recovery from assay media aliquots stored inside the incubator alongside the 24 h device experiments, normalized to immediately-frozen controls. All compounds are sorted by their hydrophobicity in terms of topological polar surface area; for abbreviations see Table 1. Data are plotted as means ± standard deviation (initial: $n$=4; 24 h: $n$=8).

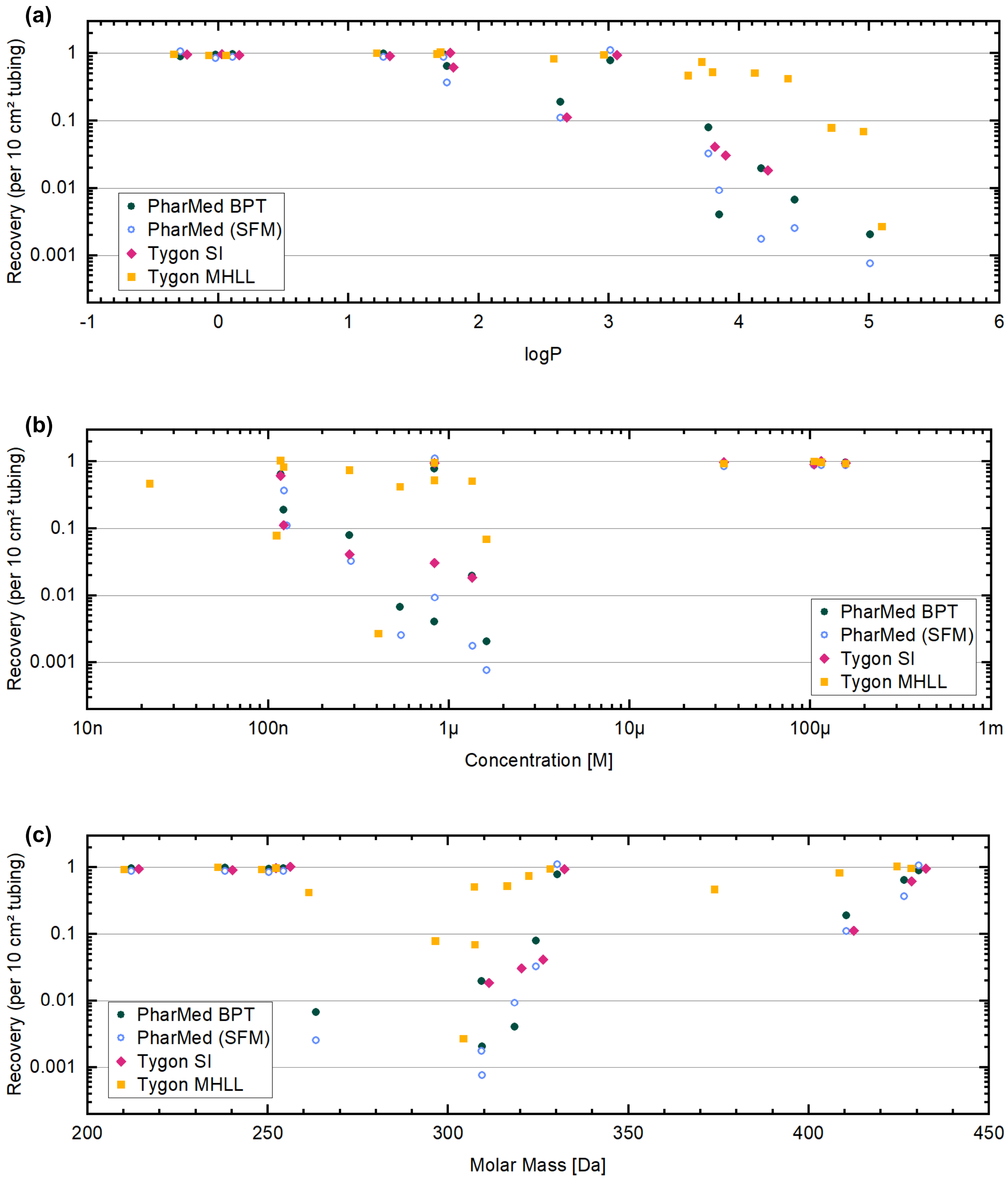


**Figure S2.** Tubing sorption as a function of compound properties. We re-plot data from Figure 3b-c as a function not of TPSA but of **(a)** log*P*, **(b)** concentration *C*, and **(c)** molar mass *M*. For clarity, we plot mean values only.

**Detailed Experimental Section**

Full details on materials and equipment can be found in the last Section.

*Microfluidics Devices*

Middle layers, identical across all device types, were prepared by cutting PC film to suitable size with a cutting plotter, adding also through-holes as seen in Fig. 1.

PDMS layers were fabricated using Sylgard 184 at 1:5 (bottom layers) or 1:20 (top layers) ratios of curing agent to base. We employed photolithographically-structured molds from SU-8 on top of silicon wafers (silanized to facilitate de-molding). Bottoms were cast at ~1 mm thickness, tops at ~3.5 mm, and cured for 60 minutes at 80 °C. We added inlet/outlet holes using a 2 mm diameter biopsy punch. PDMS layers were manually aligned with the PC film middle layer, and clamped together in a custom-made experimental rig.

OSTEmer devices were fabricated using geometrically equivalent molds (with the above) made from aluminum by CNC milling (Teflon-coated to facilitate de-molding). Unlike the pour-over PDMS molds, OSTEmer molds were designed for injection molding after enclosure with an optically-clear, non-stick film (release liner). To create inlets/outlets, and to prevent leakage, we manually added suitably-shaped PDMS pieces on top of the aluminum to create full-height inlet/oulet voids, and to serve as gaskets, respectively. After enclosure with the film, we clamped the mold stack with glass slides on top for added support, and injected liquid OSTEmer. We UV-cured the material for 180 s (bottoms; 1 mm height) or 300 s (tops; 2 mm height) at ~10 mW/cm², and then carefully delaminated OSTEmer layers from the mold. We manually aligned them with the PD film middle layer, and added gold-coated tubular rivets to the top-layer inlet/outlet holes to serve as tubing connectors. The entire stack was clamped in a custom rig, and cured at 110 °C for 72+ h.

For PC/PSA devices, we first layered PSA tapes into double-height stacks to achieve comparable 200 µm layer height. We cut channels into the layered tape using a cutting plotter, and then manually algined and bonded it to the PC film middle layer. We further bonded the tape to another PC layer to serve as the structural bottom, and to a commercial microfluidic PC connector plate to serve as the structural top. A full protocol is available on Metafluidics.[1]

*Microfluidic Setup & Tubing*

All fluidic connections for device experiments utilized PharMed BPT tubing, with stainless steel pins serving as interconnects between tubing pieces. We utilized plunger-less syringes with blunt needles as the liquid reservoirs. The liquid was guided from these into the devices, circulated through both bottom and top channels, into the peristaltic pump, and back into the reservoir. To ensure an equal total microfluidic flow length in each type of device, the inherently 2/3 shorter PC/tape device flows were supplemented with an additional half-channel (i.e., employing a flow circuit of top-bottom-top, or bottom-top-bottom). Most tubing here featured 0.25 mm diameter, whereas device connections employed 0.51 mm diameter tubing more suited to our device interfaces. At 79 cm total tubing length, the average tubing diameter in these fluidic circuits was 0.285 mm.

Tubing-only control circuits were constructed from identical-length tubing segments (including interconnects etc.), omitting only the devices themselves. For Tygon SI and Tygon MHLL we relied solely on 0.51 mm and 0.38 mm diameter tubing, respectively, based on tubing availability. We account for the altered diameters in our analysis.

*Test Solutions*

Most test chemicals were obtained as standard solutions in suitable solvents. Where this proved infeasible, we prepared stock solutions from powders in Argon-purged DMSO at the respective manufacturer-supplied solubility limits. We also used Argon-puged DMSO to pre-dilute liquid standards where needed for handling. All compounds were stored according to manufacturer guidelines before and after use. All low-volume (< 500 μl) pharmaceutical pipetting utilized low-retention tips. For intermediate handling or storage we utilized low-binding centrifuge tubes. All other plastics used in liquid handling were either polypropylene or fluorinated ethylene propylene.

We prepared a minimal media from MEM, supplemented with 0.2% Primocin and 10% serum replacement (omitted for SFM). Media for the entire experiment was prepared as a single batch. For the assay media, we added the test compounds at the concentrations from Table 1 (as well as $4.5\times10^{-5}$ g/L Zuclopenthixol and $2.7\times10^{-4}$ g/L quinine sulfate; cf. Analysis section), immediately prior to the experiment. The resulting DMSO and methanol concentrations were 1% and 0.33%, respectively. The media (with or without pharmaceuticals) was allowed to equilibrate in the incubator to 37 °C, 5% $CO_2$ for 30 minutes before being introduced to the device reservoirs.

*Experimental Procedure*

All devices and tubing were exposed to air plasma for 60 s (100 W, 1 Torr) to remove organic residues and facilitate initial wetting on day −1. After setting up fluidics as described earlier, internal surfaces were disinfected with 70% ethanol for approximately 5 minutes under flow, taking care to flush out bubbles. We subsequently rinsed fluidics with PBS (0.2% Primocin) and continued perfusion overnight at 1% pump speed inside the incubator to calibrate the pump flow rate and check for device leakage. On day 0, we flushed the fluidics with media and looped the outlet tubing into the respective inlet reservoirs for recirculating flow. In the next 24 h, we primed the fluidics by continuous recirculation of 2 ml media per device at $Q$=4.0 ml $h^{-1}$ inside the incubator (Tygon MHLL: 6.0 ml $h^{-1}$; Tygon SI: 10.8 ml $h^{-1}$).

We intermittently (4× over 24 h) manually flush the fluidics at high flow rates (>30 ml $h^{-1}$) for short intervals (<10 s) to clear bubbles. On day 1, we emptied the reservoirs and flushed the fluidics with the assay media to minimize dilution with blank media. We again set up a continuous recirculating flow over the next 24 h in the incubator, utilizing 2.5 ml assay media per fluidic circuit at the same flow rate as previously. On day 2, we collected samples for analysis from each of the reservoirs, froze them immediately to −30 °C, and moved them to −75 °C for storage within hours. For transport, samples were kept on dry ice.

*Analysis*

For fluorescence analysis, we relied on a plate reader. We were unable to detect quinine sulfate in any sample even after pH adjustment; it was thus omitted from further analysis. For the other compounds, we optimized $\lambda_{ex/em}$ as follows (with 5/10 nm bandwidths): 501/521 (F*); 551/581 (T*); 610/655 (N*). Calibration curves were created from serial dilutions of immediately-frozen day 1 stock solutions (normalized concentration $C'$=1), and corresponding "blank" media. Based on quality-of-fit ($R^2$ = 0.999), we chose a pseudo-sigmoidal correlation model of:

$$\log(Signal) = \alpha + \frac{\beta - \alpha}{1 + \left(\frac{\delta}{\log C'}\right)^{\gamma}}$$

with Greek letters denoting fit parameters. As with all statistical analysis (except where noted), we carried this out in OriginPro.

For LC/MS analysis, samples were analyzed at the clinical laboratory unit of the local university hospital using standard validated assays. The Zuclopenthixol assay proved highly unstable between different aliquots of the same sample and was thus excluded from further analysis.

Lastly, we cut tubing cross-sections using a scalpel for SEM. The low-vacuum system uses 15 kV acceleration voltage and records backscatter electron signals; the charge-up reduction mode allowed us to image samples without prior metallization. We relied on Fiji to optimize brightness and contrast.[2]

After calculating confidence intervals and means, we relied on MOVER-R (implemented in Excel) for our ratio-based analyses.[3] For partial least square analysis (PLS), we input molar concentration as $\log(C/M)$ as well as the following ChemAxon Chemicalize parameters:[4] molar mass $M$, asymmetric atom count, rotatable bond count, ring count, aromatic ring count, hetero ring count, fraction of $sp^3$ carbon atoms, H-bond donor count (H-bd), H-bond acceptor count, $\log(TPSA)$, polarizability, charge at pH 7.4, $\log P$, $\log D_{7.4}$, $\log S$, $\log S_{7.4}$, van der Waals volume, van der Waals surface area, solvent accessible surface area, minimum projection area, maximum projection area. For molecules with multiple equilibrium states for double bonds/aromatic/ring structures (i.e., the fluorophores), we utilize parameters from the dominant form at physiological pH. For some of these, $\log S$ could not be calculated, in which case we instead used the average (across analogs) as the model input.

#### Materials & Equipment

##### *Neuropsychopharmaca & Fluorophores*

| Product | Manufacturer | Identifier |
|---|---|---|
| Amitriptyline (Am) 1 mg/ml in Methanol | Cerilliant | A-923 |
| Citalopram (Ci) 0.1 mg/ml in Methanol | Cerilliant | C-057 |
| Clomipramine (Cl) 1 mg/ml in Methanol | Cerilliant | C-118 |
| Fluoxetine (Fl) 1 mg/ml in Methanol | Cerilliant | F-918 |
| Haloperidol (Ha) 1 mg/ml in Methanol | Cerilliant | H-030 |
| Lacosamide (La) powder | European Pharmacopoeia (EP) | Y0001982 |
| Licarbazepine (Li) powder | Cayman Chemical | 18467 |
| Methadone (Me) 1 mg/ml in Methanol | Cerilliant | M-007 |
| Nortriptyline (No) 1 mg/ml in Methanol | Cerilliant | N-907 |
| Paliperidone (Pa) 1 mg/ml in Methanol | Cerilliant | H-076 |
| Risperidone (Ri) 1 mg/ml in Methanol | Cerilliant | R-006 |
| Rufinamide (Ru) powder | Cayman Chemical | 18870 |
| Sertraline (Se) 1 mg/ml in Methanol | Cerilliant | S-021 |
| Vortioxetine (Vo) powder | Cayman Chemical | 30183 |
| Zonisamide (Zo) powder | Cayman Chemical | 24183 |
| Zuclopenthixol powder | Cayman Chemical | 24961 |
| Reference Dye Sampler Kit (1mM; includes quinine sulfate, fluorescein, carboxy-TRITC, sulforhodamine 101, nile blue) | Invitrogen | R14782 |

##### *Other Reagents*

| Product | Manufacturer | Identifier |
|---|---|---|
| Dimethyl sulfoxide | Sigma Aldrich | D2650 |
| DPBS, − calcium, − magnesium | Gibco | 14190144 |
| Minimum Essential Medium (MEM), no glutamine, no phenol red | Gibco | 51200046 |
| KnockOut Serum Replacement | Gibco | 10828010 |
| Primocin | Invivogen | ant-pm |

##### *Microfluidics*

| Product | Manufacturer | Identifier |
|---|---|---|
| PDMS Silicone Elastomer Kit | DOW | Sylgard 184 |
| OSTE+ Crystal Clear | Mercene Labs | OSTEMER 322 |
| OSTE+ Release Liner | 3M | 9742 |
| Tubular rivets, gold plated (DIN 7340 A 2.5×0.3×3.5 mm) | Kaiser Waltermann | |
| Double Sided Medical Tape | 3M | 9877 |
| Microfluidic interface (polycarbonate, microscopy slide format, 2x16 olives) | Microfluidic chipshop | 10-1121-0343-03 |
| Polycarbonate film (125 µm) | Covestro | Makrofol DE 1-1 |
| Syringes, 5ml (as liquid reservoirs) | Restek | 22774 |
| Blunt dispensing needles 23G | Metcal | 923050-TE |
| Steel Microfluidic Fittings 23G | Elveflow | LVF-KFI-13 |
| PharMed BPT peristaltic pump tubing | Ismatec | SC0320 |

| | | |
|---|---|---|
| PharMed extension tubing | Ismatec | SC0337 |
| PharMed extension tubing | Ismatec | SC0339 |
| Tygon MHLL peristaltic pump tubing | Ismatec | SC0716 |
| Tygon SI peristaltic pump tubing | Ismatec | SC0620 |

*Equipment*

| **Product** | **Manufacturer** | **Identifier** |
|---|---|---|
| Low-pressure plasma system | Diener | Femto |
| Cutting Plotter | Graphtec | CE5000 |
| 16-channel peristaltic pump with click-‘n-go cartridges | Ismatec | IPC-N |
| $CO_2$ incubator | Thermo Scientific | Heracell Vios 160i |
| Multimode Plate Reader | Tecan | Infinite M1000 Pro |
| Tabletop SEM | Hitachi | TM-1000 |
